# Pilot scale study on UV-C inactivation of bacterial endospores and virus particles in whole milk: evaluation of system efficiency and product quality

**DOI:** 10.1101/2022.01.07.475436

**Authors:** Pranav Vashisht, Brahmaiah Pendyala, Ankit Patras, Vybhav Vipul Sudhir Gopisetty, Ramasamy Ravi

**Affiliations:** Food Biosciences and Technology Program, Department of Agricultural and Environmental Sciences, Tennessee State University, Nashville, 37209, TN, USA

**Keywords:** Pilot scale UV system, whole milk, bacterial endospores, virus particles, lipid peroxidation, volatiles profile

## Abstract

UV-C processing of whole milk (WM) using a designed pilot scale Dean flow system was conducted at flow rates (11.88, 23.77, and 47.55 gph), Reynolds number ranges from 2890-11562 and the Dean number (at curved region) calculated as (648-2595) to inactivate bacterial endospores and virus particles. Biodosimetry studies were conducted to quantify the reduction equivalent fluence at selected experimental conditions. Results revealed that the fluence distribution improved as flow rate increases, attributed to increase in Dean effects and turbulence intensity. Microbial inactivation studies conducted at 47.55 gph showed 0.91 ± 0.15 and 2.14 ± 0.19 log reduction/ pass for *B. cereus* endospores and T1UV phage. Linear inactivation trend was observed against number of passes which clearly demonstrates equivalent fluence delivery during each pass. Lipid peroxidation value and volatiles profile did not change significantly at UV fluence of 60 mJ/cm^2^. Lower E_EO_ value signifies the higher electrical efficiency of the system.

## 1. Introduction

Milk and its products are the potential source of various nutrients such as lipids, proteins, carbohydrates, vitamins, minerals, and enzymes; these components present in milk allows the growth of numerous pathogenic microorganisms (Oliver et al., 2005; Shanmugam et al., 2012; Bogahawaththa et al., 2018; Munir et al., 2019). According to Delorme et al. (2020), dairy processing needs strict quality control as contamination issues can originate at any stage of the food chain. The presence of aerobic spore-forming bacteria of *the Bacillus* genus in milk is consider as the major cause of deterioration of raw and pasteurized milk (Crielly et al., 1994; Cosentino et al., 1997). *B. cereus* is a major concern for the dairy industry because of its psychrotolerant nature, which limits the shelf life of the pasteurized milk and milk products stored at 6-7°C and its ability to produce toxins which makes it a potential cause for foodborne illnesses where the production of emetic and enterotoxin results in vomiting and diarrhea, respectively (Griffiths, 1992; Bilbao-Sáinz et al., 2009). According to CDC, more than 63000 cases of *B. cereus* illness are reported annually in the United States and all of them are foodborne (Scallan et al., 2011).

Pasteurization and sterilization are the most common techniques used by industries to ensure the safety of milk (Bermúdez-Aguirre et al., 2009; Choudhary and Bandla, 2012; Patras et al., 2020). The Code of Federal Regulations (3), Title 21, Section 131.3b, defines pasteurized dairy products as using properly operated equipment to heat every particle of the product to a specified temperature and held at or above that temperature for a specified time. Though these techniques are efficient and can inactivate numerous foodborne pathogens ranging from vegetative cells to viruses and spoilage causing enzymes. One problem that is found to cause a reduction in the shelf life of WM is the presence and growth of *Bacillus* sp. and its heat resistant spores in pasteurized, sterilized as well as ultra-pasteurized milk (Pettersson et al., 1996; Scheldeman et al., 2004; Cullor, 2011). The high pasteurization temperatures can easily destroy vegetative cells but can activate the spores, which leads to germination and growth (McGUIGGAN et al., 1994). Other disadvantages of pasteurization techniques are significant alteration in the sensorial and nutritional profile of the product, protein denaturation, undesired changes to milk fat globules, high operational and processing cost which makes it unfeasible for small scale dairy units (Garcia-Amezquita et al., 2009; Cappozzo et al., 2015; Gunter-Ward et al., 2018). Raw dairy products are prioritized by the consumers because of their typical sensorial characteristics and other health benefits such as decreased incidence of asthma, allergies, and respiratory infections which suggest immune benefits of dairy proteins are lost upon heat processing (Buchin et al.,1998; Braun-Fahrländer and Von Mutius, 2011). To preserve the benefits of raw dairy products without compromising the safety warrants the study of non-thermal technology for processing of dairy products (Gunter-ward et al., 2018). Interest in light-based technologies (UV-C irradiation) for liquid food processing has exponentially grown over the years is increasing due to lower energy consumption, cost-effectiveness, minimal effect on the quality of the products (Tran and Farid, 2004; Keyser et al., 2008; Corrales et al., 2012; Feng et al., 2013; Ochoa-Velasco et al., 2014; Jermann et al., 2015; Islam et al., 2016). Several federal agencies have approved the use of UV-C technology for milk treatment. For example, European Food Safety Authority (EFSA) approved the use of UV-C at 253.7 nm for milk processing post pasteurization for the extension of shelf life and to improve the nutritional value as it increases the vitamin D3 content in the product (EFSA, 2016, and Koutchma, 2018). Food Safety and Standards Authority of India approved the UV-C processing of raw milk processed via the Sure Pure UV system (Koutchma, 2018). In USA, any new technology for processing of grade A milk has to go through PMO and FDA clearance.

Many lab-scale studies on the UV-C processing of milk revealed the efficiency of the process (Choudhary et al., 2011; Bandla et al., 2012; Gunter-ward et al., 2018; Ansari et al., 2019). Still, the major challenges at the pilot scale are limited light penetration and mass transfer (Sizer and Balasubramanian, 1999) and fluence delivery per single pass. Milk is an opaque fluid with high absorption as well as scattering coefficient due to the presence of lipids, proteins and vitamins. UV light is absorbed by these micro and macro nutrients and creates UV gradients (Walstra and Jenness, 1984). Some studies reported that the major problem of UV-C processing is oxidation of fats and off-flavors (Bekbölet, 1990; Matak et al., 2005, 2007). However, the use of an efficient reactor design can improve mass transfer and mix the bulk field homogenously overcoming those challenges, delivering a uniform UV fluence (Koutchma, 2009). It is quite evident in the literature that Dean flow reactor is an efficient design where additional turbulence is generated when fluid passes through the coiled tube (Dean, 1927; Gopisetty et al., 2018; Vashisht et al., 2021). This is known as the Dean effect, and it is produced by the curvature radius between the inner and outer boundary layer (Dean, 1927). In our previous study we used a simulated fluid and scientifically proved that the Dean flow UV reactor distributes the fluence homogeneously demonstrated by the linear microbial inactivation kinetics (Vashisht et al., 2021). In reactor engineering, it is crucial to ensure that all the fluid particles receive the UV fluence within the acceptable limit i.e. minimal product damage and efficient microbial inactivation. An understanding of the fluid mechanical and flow properties is extremely important. Herein, we will evaluate the peroxidation and volatiles of WM, confirming the UV fluence (pasteurization equivalent dose). We hypothesize that effective fluence delivery in the system can inactivate the pathogens and preserve the quality of the product. The objectives of the present study are i) validation of the pilot-scale Dean flow UV system using biodosimetry approach, ii) evaluation of the microbial inactivation kinetics, iii) assessment of the volatile profile and lipid peroxidation of the UV irradiated whole milk.

## 2. Materials and Methods

### 2.1 Bacteriophage and endospores propagation

T1UV bacteriophage and *Bacillus cereus* ATCC 14579 endospores were used in this study. T1 UV culture (@10^10^ pfu/mL) was purchased from GAP EnviroMicrobial Services Limited, London, ON, Canada and *Bacillus cereus* ATCC 14579 was obtained from American Type Culture Collection (ATCC) center (Manassas, VA). Culturing of endospores was conducting using a Bacto Brain Heart Infusion Broth (BHI, Beckton, Dickinson, Franklin Lakes, NJ) as a growth medium, incubation was done at 37 °C with aeration at 180 rpm. Nutrient-rich, a chemically defined medium known as Mineral Salts Medium (MSM) was used for sporulation. 20 mL of overnight grown culture was inoculated in 200 mL of MSM in 1 L of the flask. Incubation was be done at 30 °C with aeration at 180 rpm for 3 days. Dormant spores were purified by suspending the pellets in 20% Nycodenz (VWR, Atlanta, GA) followed by 50% Nycodenz density gradient centrifugation at 14000 g for 45 minutes. Enumeration was done using a standard plate count method.

### 2.2 Optical properties

Optical properties (absorbance and reduced scattering) of the WM were evaluated at 254 nm by using a double beam spectrophotometer connected to a 6-inch integrating sphere. Inverse addling doubling technique was used to quantify the absorption coefficient, scattering coefficient, base-e. A series of thin quartz cuvettes were used to achieve a light transmission between 10 to 14%. A MS-DOS based code was run to separate the absorption coefficient and reduced scattering coefficients. Light absorption and reflection were accounted for in the calculations.

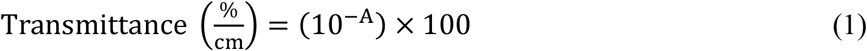

where A stands for the absorbance (base 10) of the sample at 254 nm for a 1 cm path length.pH of the product was measured by using a pH meter (Jenway, Cole Palmer, OSA, UK). All the measurements were taken in triplicate to reduce error.

### 2.3 Organism sensitivity test

*Bacillus cereus* ATCC 14,579 and T1UV phage were evaluated for D_10_ values (dose required for 90 % inactivation or 10 % survival). Microorganisms were inoculated separately (@10^6^-10^7^ CFU/mL or PFU/mL) in whole milk. The absorption coefficient of the final solution was measured using the UV–Vis. spectrophotometer (Thermo Scientific, Genesys 10S, Milwaukee, WI, US). Known UV fluence were delivered (3, 6, and 9 mJ/cm^2^) for both the microorganisms (Bolton and Linden, 2003). 5 mL of sample was exposed to UV using a Collimated Beam Apparatus (Trojan Technologies, London, ON, Canada), which contains a 25 W low-pressure mercury lamp. The suspension was stirred during irradiation using a magnetic stirrer to ensure equal UV volatil to all microorganisms. The D_10_ value was evaluated using the LINEST function of Microsoft Excel between the UV fluence delivered, and the log inactivation obtained. All the experiments were done in triplicate to reduce the error

### 2.4 UV-C Treatment

WM was exposed to UV-C light using the Dean flow UV system (Fig. 1). The system is a continuous UV reactor, consisting of a 500W low-pressure mercury lamp protected with stainless steel cylindrical blocker, a cooling fan, and a food-grade FEP (Fluorinated Ethylene Propylene) tube. The tube length is 26 m with an inner diameter (D_i_) of 0.64 cm, an outer diameter (D_o_) of 0.74 cm, a wall thickness of 0.5 mm. The tube was wrapped around the centrally positioned lamp in a serpentine path. The diameter of the coil (D_c_) was 12.7 cm. The tubing was carefully engineered to have significant curves (bends) to induce mixing. The system also included a 4–20 mA current loop receiver and an indicator for output power lamp display. The electric supply of 220 V (single phase), 60 Hz was used. The irradiance at the surface of the FEP tube was 22 mW/cm^2^, measured using an International Light Technologies (Peabody, MA) IL-1700 radiometer with a SED 240 detector fitted with a NS254 filter. Transmittance (UVT%) of FEP tube was 65%, measured using a UV–Visible Spectrophotometer (Thermo Scientific, Genesys 10S, Milwaukee, WI, US). Therefore, the calculated UV intensity at the surface of the fluid was 14.3 mW/cm^2^. The total reactor volume was 0.80 L. The system was connected to an inlet, outlet, and a rinse tank (20 L each). The output power at 254 nm was approximately 220 Watts at 254 nm. WM was treated at a flow rate of 0.75, 1.5, and 3 L/min (equivalent to 11.88, 23.77, 47.55 gph, respectively). The flowrate was controlled by Watson Marlow 730U peristaltic pump (Watson Marlow, Charlotte, NC, US). The pump was validated by using a bucket test in which time to collect a known volume of the fluid was estimated. CIP was done before starting the experiment as well as at the end. All experiments were conducted in triplicated to reduce random error. Dean number (De) evaluated as a function of Reynolds number (Re) and the geometric parameters D and Dc, which stands for the hydraulic diameter of the tube (m) and diameter of the coil (m), respectively (Gopisetty et al., 2018, 2019). The following equation was used:

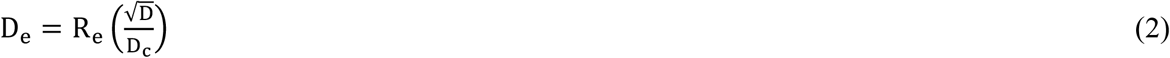

**Fig. 1.**
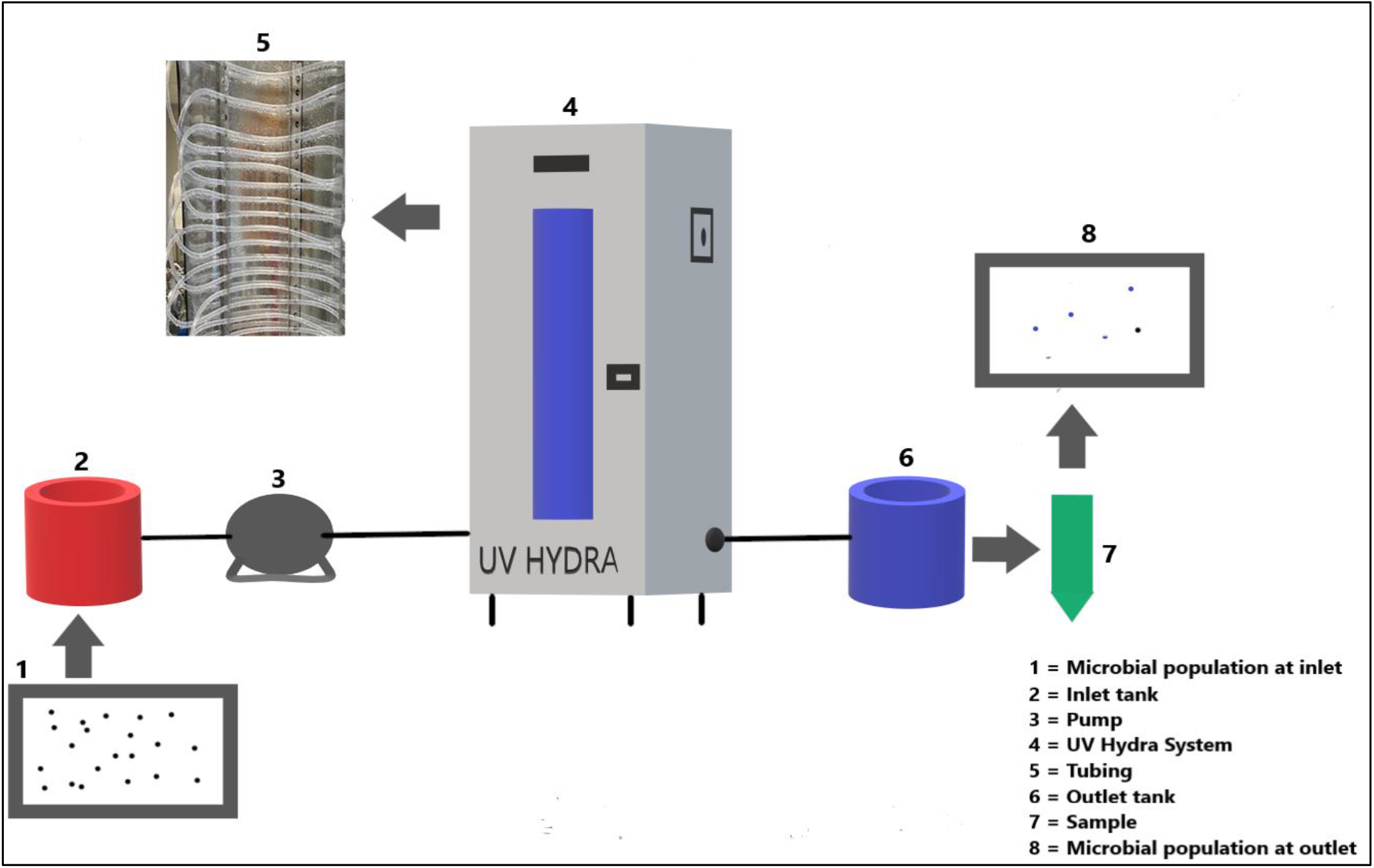
Experimental setup of Dean flow UV system.

Where Reynolds numbers (R_e_) was evaluated as:

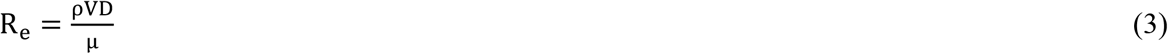

where ρ denotes the fluid density, μ denotes the dynamic viscosity, V is the fluid velocity, and D denotes the diameter of the tube. The R_e_ of < 2100 represents laminar flow pattern, R_e_ between 2100-4000 shows transient flow conditions, and R_e_ of > 4000 means turbulent flow conditions (Bandla et al., 2012; Pendyala et al., 2021; Vashisht et al., 2021). The residence time (RT) was 64, 32 and 16 seconds for 0.75 L/min. (11.88 gph), 1.5 L/min. (23.77 gph) and 3 L/min. (47.55 gph) respectively, which was evaluated by using the following equation:

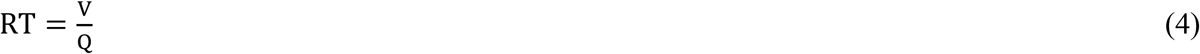

Where Vis the volume (L); Q is the flow rate (L/min) Average flow velocity (vf) (m/sec.) was evaluated by using the following equation:

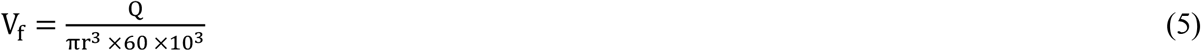

Q is the flow rate of fluid in L/min. r is the tube’s radius in m.

### 2.5. Determination of volatiles using electronic nose

Volatiles were evaluated by using an electronic nose (Heracles II, Alpha MOS, Toulouse, France). It consists of two columns working in parallel mode: a non-polar column (MXT5: 5% diphenyl, 95% methylpolysiloxane, 10 m length and 180 μm diameter). The volatiles profile of treated samples (at 60 and 120 mJ/cm^2^ of fluence) was compared with controlled (untreated) samples. 20 mL of vials were filled with 10 mL of the sample followed by sealing with septa screw cap using a crimper. The vials were equilibrated for 200 seconds at 50 °C in agitation chamber. Autosampler was used to inject the generated headspace aroma into the electronic nose column at a flow rate of 270μL/s. The column temperature program was 40°C (1 min) – 2 °C min^-1^ – 200 °C (3 min). The temperature of injector and detector was set at 180°C and 220°C, respectively. The chromatograms were obtained using Flame ionization detector (FID) and individual volatile profile including Kovats index, retention time and sensory description was evaluated by using AromaChemBase software. The analysis was done by using Alpha Soft (Alpha Soft version 3.0.0, Toulouse, France). All the samples were evaluated in triplicate to reduce error and all the values were mentioned in terms of relative area.

### 2.6. Determination of lipid peroxidation

Lipid peroxidation value was evaluated by a standard curve method using spectrophotometer. According, to Matak et al. (2007) value is measured in terms of MDA (malondialdehyde) and other reactive substance so standards of 50, 25, 12.5, 6.25, 3.125 μM concentrations were prepared using MDA. 1 mL of irradiated whole milk was mixed and vortexed with 7 mL of 1% phosphoric acid followed by the addition of 2 mL of 42 mmol/L TBA (thiobarbituric acid). The solution was heated at 100°C for 60 minutes followed by cooling it down (by keeping on ice). The color was changed to pink, where the intensity was dependent upon the extent of lipid peroxidation. 4 mL of solution was added to the 4 mL of 1:12 mixture of 2 mol/L sodium hydroxide and methanol, it was vortex mixed for 10 seconds followed by centrifugation at 13000 g for 3 minutes. Supernatant was evaluated for absorbance at 532 nm using a UV-Vis spectrophotometer (Thermo Scientific, Genesys 10S, Milwaukee, WI, US) calibrated against blank.

### 2.7. Electrical energy per order (E_EO_)

E_EO_ was estimated to measure the kilowatt-hours of energy required to reduce the microbial population by one log (Bolton, 2010). This term is commonly used to assess electrical energy efficiency (Ward et al., 2019). E_EO_ value depends upon the nature of fluid under processing and the geometry of the reactor (Pendyala et al., 2021). Evaluation was done by using the equation given by Ward et al. (2019).

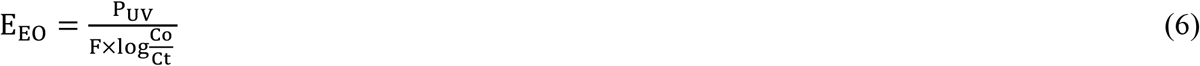

where P_UV_ refers to the power of the electric lamp or total energy of the lamp and power supply (kW), F stands for volumetric flow rate (m^3^/h.), and Co and Ct stand for initial and final microbial concentration, respectively.

### 2.8. Statistical Analysis

All the treatments were conducted in replicates where each sample was independent and randomly assigned. All the values are reported as mean ± std. deviation. For evaluation of effect of numerous treatments on REF, inactivation kinetics, lipid peroxidation and volatiles profile one-way ANOVA along with the Tukey HSD (honestly significant difference) at p<0.05 significance level.

## 3. Result and Discussion

### 3.1. Optical properties of test fluid

Optical properties of WM are illustrated in Table 1. The absorption and reduced scattering coefficient were measured as 23.7 ± 0.3 cm^-1^ (transmittance - 2.5E-22 %/cm) and 42.3 ± 0.5 cm^-1^ respectively, with 12 % reflectance. The pH and fat content of the WM was 6.73 ± 0.17 and 3.27 %, respectively. Previous studies evaluated the absorption coefficient of whole milk without separating the reduced scattering coefficient and reported significantly higher values, 220 cm^-1^ (Choudhary et al., 2011), 326 ± 1.5 cm^-1^ (Ansari et al., 2019). Optical properties clearly show the opaque nature of the test fluid. WM significantly absorbs and scatters the UV photons. Suspended particles are mainly responsible for attenuation of UV light via scattering effect (Gunter-ward et al., 2018). If the product particles are larger than the UV wavelength then it leads to the forward scattering with amplified back scattering effect (Koutchma, 2009). Our results demonstrate high scattering effect of WM, due to fat globules and proteins (micellar casein) as reported in the literature (Gunter-ward et al., 2018; Ward et al., 2019). There could be a minor scattering effect due to dissolved minerals and sugar (lactose), however it can be ignored because their proportionate size is lesser than the fat and protein components (Gastélum-Barrios et al., 2020).

**Table 1.**
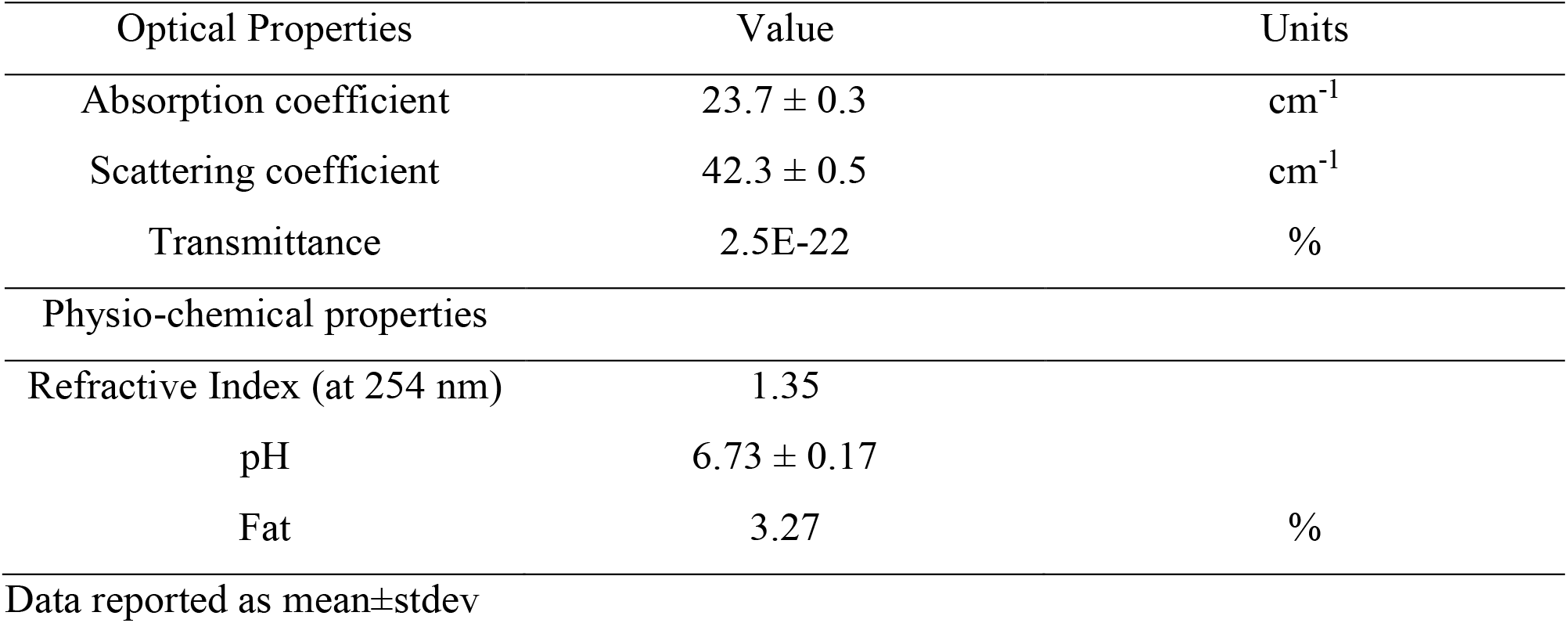
Optical and Physio-chemical properties of whole milk

### 3.2. Design and Development of dean flow reactor

Many literature studies used circular path coiled tube dean flow UV systems to evaluate microbial inactivation in milk (Choudhary et al., 2011; Bandla et al., 2012; Ansari et al., 2019). To improve the Dean forces, in our previous proof-of-concept study we designed and developed pilot scale serpentine path coiled Dean flow UV system and evaluated its microbial inactivation efficiency. Previously, we irradiated fluids with absorption coefficient ranging from 6.5 to 17 cm^- 1^ at commercial flow rates (Vashisht et al., 2021) and evaluated the system performance. In the current study, the system was challenged against a highly opaque fluid such a WM with an absorption coefficient of 23.7 cm^-1^. The initial design of the system consisted of a Teflon tubing (45% UV transmission) of 1 cm path length, 20 m total length and 18 cm coil diameter (Vashisht et al., 2021). The reactor system was modified where FEP tubing (65% UV transmission) was used to have high transmittivity. Tube length was increased to 26 m to increase the residence time and UV fluence/pass. The optical thickness of the fluid was reduced to 0.64 cm. This will allow higher penetration of photons in the bulk fluid. The coil diameter was reduced to 12.7 cm to increase the Dean effect (as Dean number is inversely proportional to coil diameter). The ratio of D to Dc (where D stands for inner diameter of tubing (0.64 cm), and Dc stands for the diameter of coil (12.7 cm)) of the designed reactor was 0.050 (0.64/12.7), which was in the range of 0.03–0.10 demonstrating the initiation of Dean flow vortices during the experiments (Koutchma, 2009). Bandla et al. (2012) used Reynolds number as a parameter of turbulence but missed to evaluate the Dean number. This number is a significant parameter as it generates the additional turbulence hence the efficient mixing conditions (Dean 1927, Gopisetty et al., 2018, 2019). The evaluated Reynolds and Dean number for each flow rate was in the range of 2890-11562 and 648-2595, respectively (Table 2). This data revealed that the experiments were conducted at transient to fully turbulent conditions to provide efficient mass transfer (Dean, 1927; Patras et al., 2020). Turbulent flow conditions deliver higher velocity to the particles as compared to the mean velocity of individual particles providing efficient UV fluence distribution (Koutchma, 2009). Volume average UV intensity depends on the relative positions of the particle and the UV lamps orientation. The UV fluence received by the micro-organisms depends on two main factors: the trajectories through the system and the REF rate (Vashisht et al., 2021). The incident radiation rate is dependent on the positions of the UV lamps within the UV system.

**Table 2.** Flow properties of the WM under testing

| Flow rate<br>(L/min.) | Residence time<br>(sec.) | Velocity of flow<br>(m/sec.) | Reynolds number | Dean number |
| --- | --- | --- | --- | --- |
| 0.75 | 64 | 0.39 | 2890 | 648 |
| 1.5 | 32 | 0.78 | 5781 | 1297 |
| 3 | 16 | 1.55 | 11562 | 2595 |

### 3.3. UV fluence validation at experimental flow rates

Biodosimetry study was conducted using *B. cereus* endospores to quantify delivered UV fluence. The D_10_ value of *B. cereus* endospore was quantified using a standard collimated beam system as 8.91 ± 0.07 mJ/cm^2^, it was within the range as reported in the literature (Gopisetty et al., 2018; Pendyala et al., 2019, 2020, 2021). WM was inoculated with *B. cereus* (10^7^ cfu/mL) and it was passed through the UV systemlight at flow-rates of 0.75 L/min. (11.88 gph), 1.5 L/min. (23.77 gph) and 3 L/min. (47.55 gph). REF was evaluated as a product of D_10_ value and log reduction (LogN_0_-N). Total REF was 26 ± 1.07, 14.5 ± 0.53 and 10 ± 0.27 mJ/cm^2^ at 11.88, 23.77 and 47.55 gph of flow rate, respectively. It was observed that with the increase in flow rate, total REF was significantly reduced due to lower hydraulic retention times (Fig. 2a). In contrast, the REF rate (REF per sec.) was observed to be increase with the increase in flow rate attributed to the increase in turbulence and Dean forces intensity as demonstrated by the increased Reynolds and Dean number (Fig. 2b). Similar trends were reported by previous authors (Gunter-Ward et al., 2018; Ansari et al., 2019; Vashisht et al., 2021). According to Vashisht et al. (2021) Dean number plays a pivotal role in UV fluence distribution while the residence time directs the total REF for the inactivation of microbes hence a balance needs to be maintained between these process parameters. Herein, REF rate can be a useful factor for the determination of system’s performance in terms of fluence distribution which is critical at commercial level. At 47.55 gph flow rate, the REF rate was 1.54 and 1.38 times efficient than the 11.88 and 23.77 gph of flow rate. UV-C systems are suitable for the treatment of opaque liquids when efficient mixing conditions that can surpass the potent intensity gradient and ensure effective fluence distribution (Vashisht et al., 2021). Inefficient mixing conditions will significantly affect the UV system performance and will result in non-linear inactivation kinetics of the target pathogen (Atilgan, 2013; Crook et al., 2015). Completing understanding of the UV system performance, knowledge of optical properties of the test fluids will allow accurate sizing of UV systems.

**Fig. 2.**
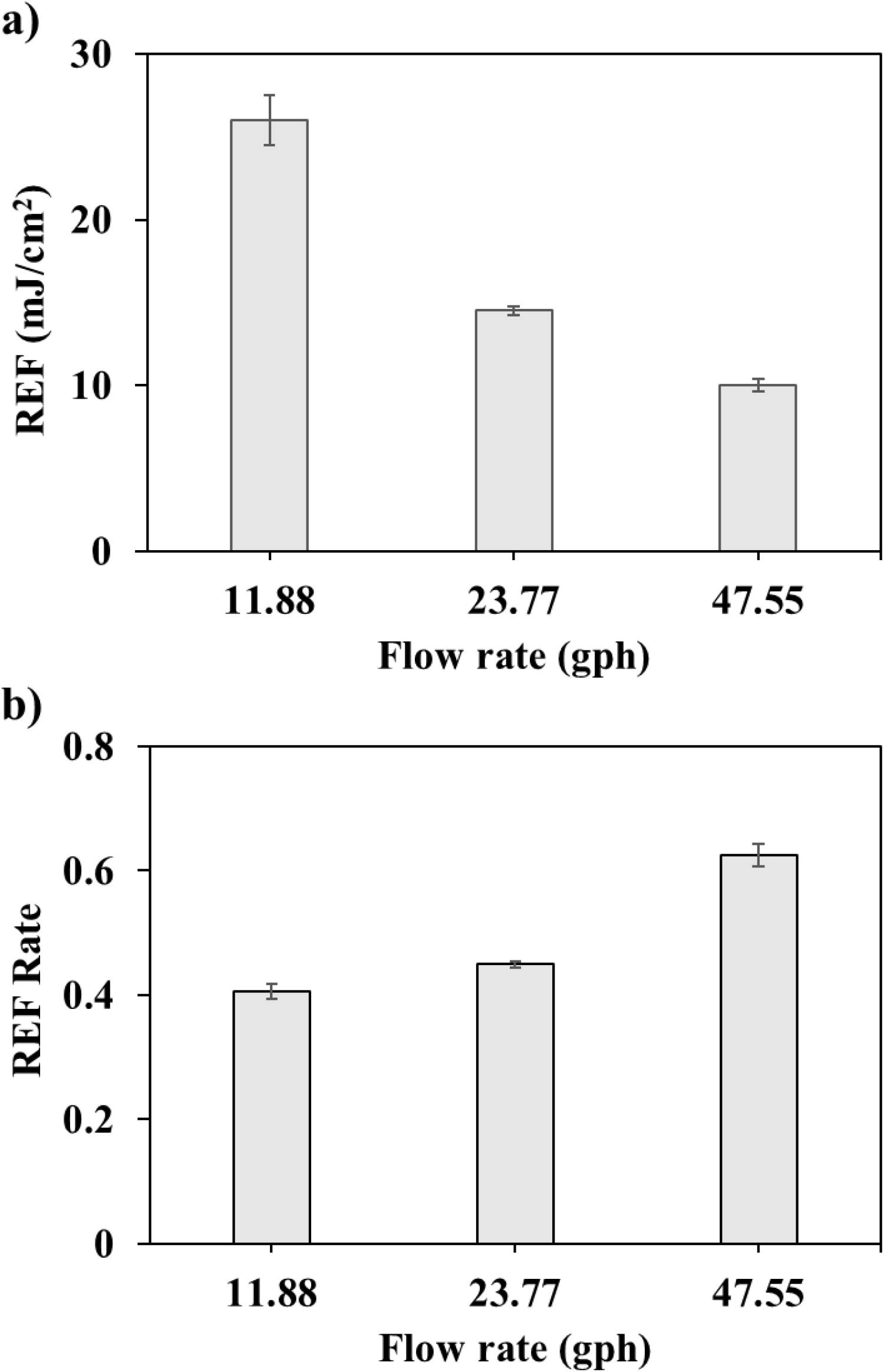
Effect of flow rate on delivered fluence (bio studies) a) REF b) REF rate Note: REF rate = REF/residence time

### 3.4. Microbial Inactivation studies

Since better system performance was observed at higher flow rate, microbial inactivation studies were conducted at a flowrate of 47.55 gph using *B. cereus* endospores and T1UV virus particles (selected as a surrogate to *Salmonella, Listeria* and *Cronobacter*). Selection of pertinent microorganism is of utmost importance during validation studies (Patras et al., 2020). *B. cereus* endospores are most common pertinent bacterial spores present in milk and causes food safety and spoilage issues (Choudhary et al., 2011, Gunter-Ward et al., 2018; Patras et al., 2020) while *Salmonella & Listeria* are the pertinent vegetative microorganisms in milk (Patras et al., 2020). The D_10_ value of T1UV phage is similar to *Salmonella* and Listeria and therefore used in this research study. 0.91 ± 0.15 and 2.14 ± 0.19 log reduction per pass (R^2^ > 0.99, p < 0.05) was observed for *B. cereus* and T1UV phage, respectively (Fig. 3a and 3b). Linear trend demonstrates uniform REF delivery during each pass. The kinetics of the UV inactivation are widely assumed for a first-order reactions with respect to the REF. The typical UV response curve can be represented in two ways: a log survival curve and a log inactivation curve

**Fig. 3.**
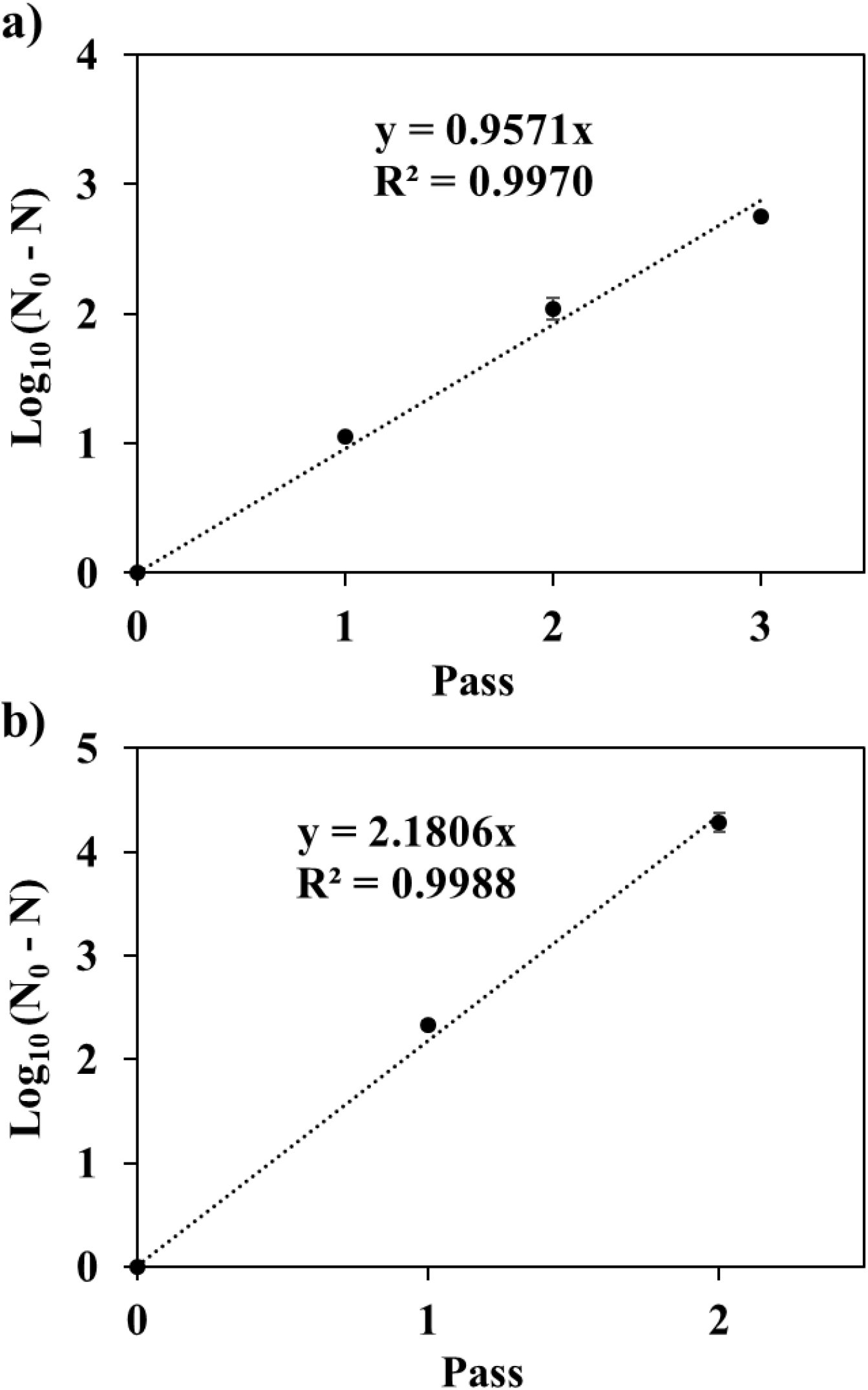
Microbial inactivation per each pass of WM a) *B. cereus* endospores b) T1UV phage

It is quite evident if the system is not efficient and does not provide good mixing conditions, then the desired UV fluence will not be delivered to the microbes which may result in lower inactivation rate at higher UV fluence (Bhullar et al., 2019) and tail may be observed in inactivation curve. For instance, Makarapong et al. (2020) reported 4.7 and 4.6 log reduction of total microbial count in WM treated with 98 and 110 mJ/cm^2^ of UV fluence. In our studies total log reduction of *B. cereus* and T1 UV phage were 2.75± 0.01 and 4.29 ± 0.09 after three and two passes, respectively. Previous Dean concept studies reported a significant log reduction of pathogenic microbes in dairy products. Ansari et al. (2019) reported *B. cereus* endospores log reduction of 6, 2.90 and 1.1 in skim, cow and bovine milk, respectively where UV-C processing was used as hurdle technology. Bandla et al. (2012), reported 2.3 log reduction of standard plate count in raw cow milk given a UV-C fluence of 16.82 mJ/cm^2^. Matak et al. (2005) reported 5-log reduction of *Listeria monocytogenes* in goat milk given a UV-C fluence of 15.8 ± 1.6 mJ/cm^2^ but the inactivation kinetics reported was non-linear which demonstrates that author failed to achieve efficient mixing conditions. Crook et al. (2015) used thin film reactor and reported 5-log reduction of *Listeria monocytogenes* and *Staphylococcus aureus* in WM, but the fluence was reported in J/L which is disparate to the commonly used standard unit i.e., mJ/cm^2^. In our previous studies we introduced the concept of Pasteurization and Sterilization Equivalent Dose referred to the fluence required for 5 log reduction of *S*. Typhimurium and 6 log reduction of *B. cereus* endospores, respectively (Vashisht et al., 2021). Therefore, in our studies at 47.55 gph of flow rate, 3 and 7 passes are sufficient to provide a pasteurization and sterilization equivalent dose, respectively while another approach could be scaling up and increasing the length of tubing of the present system to increase the residence time and hence the delivered fluence per pass.

### 3.5. Lipid peroxidation

Milk consists of photosensitizers compounds such as riboflavin which can cause lipid peroxidation during UV-C light exposures and can deteriorate the quality and limits the shelf life of the dairy products (Bekbolet, 1990). The product of these reactions are hydroperoxides that can decompose and form free-radicals which further causes auto oxidation and formation of off-flavoring compounds (Bekbolet, 1990). The oxidation reactions are prompt and usually take place when milk is exposed to metal surface or light (van Aardt et al., 2005). According to Cadwallader and Howard, 1998 autoxidation and light induces oxidation of unsaturated fatty acids of milk are primarily responsible for the origination of off flavor and reduction in nutritive compounds like ascorbic acid and riboflavin. Literature studies reported that the extent of lipid peroxidation is directly proportional to riboflavin content, and it is significantly affected by the exposure to UV light photons (van Aardt et al., 2005). In our studies the pertinent microorganism was *B. cereus*. Hence, sterilization equivalent dose for *B. cereus* was 53.46 mJ/cm^2^ (6 × 8.91 mJ/cm^2^) (Vashisht et al., 2021). Therefore 60 mJ/cm^2^, 120 mJ/cm^2^ and 180 mJ/cm^2^ of UV fluence were selected for lipid peroxidation studies. The temperature during the UV light exposures processing was maintained at 4 ° C to nullify the effect of lipase activity on fatty acids in WM (Deeth and Fitz-Gerald, 1995). Fig. 4 demonstrates the increase of lipid peroxidation in a polynomial trend (2 order equation) with the increase in the delivered UV fluence. Controlled samples had positive MDA value (0.110 ± 0.002 μM), which shows that the lipid peroxidation was initially present in the WM. At UV-C fluence of 60, 120 and 180 mJ/cm^2^, MDA concentration was 0.114 ± 0.004, 0.138 ± 0.005, and 0.168 ± 0.002 μM, respectively. Fig 4b demonstrates that there was no significant difference (p<0.05) between the MDA concentration of the controlled samples and the samples treated with UV-C fluence of 60 mJ/cm^2^, enough to inactivate 6.73-log of *B. cereus* and 12.58-log of T1 phage. Makarapong et al. (2020) reported a significant lipid peroxidation in WM samples treated with 98 and 110 mJ/cm^2^ of fluence. However, the evaluated fluence was significantly higher for the pasteurization equivalent dose reported in this study.

**Fig. 4.**
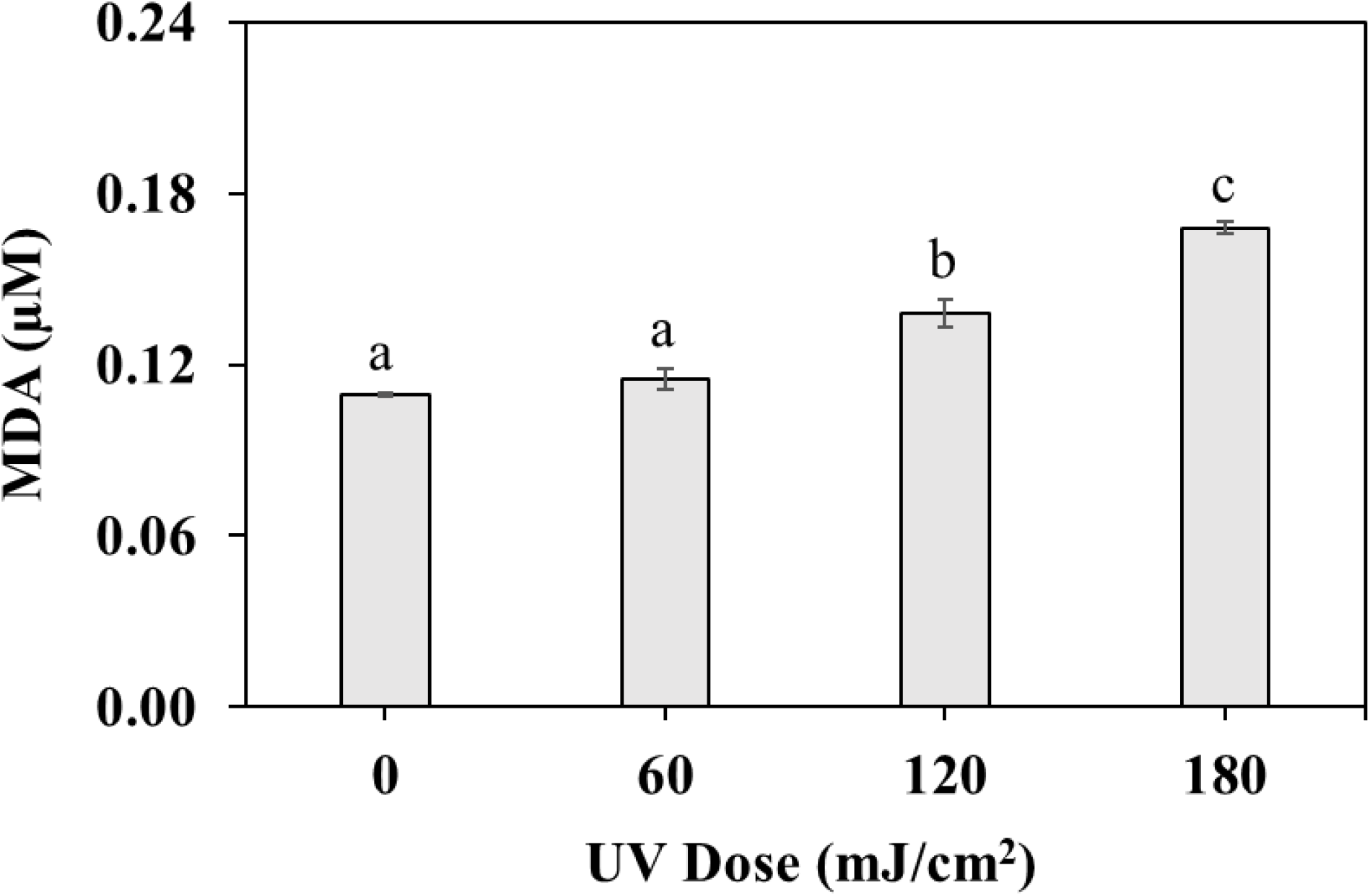
Variation of malondialdehyde concentration in irradiated milk samples with respect to the delivered UV fluence.

### 3.6. Volatiles Profile

Table 3 represents the volatiles profile of the untreated and irradiated WM samples. The irradiated samples were exposed to UV fluence of 60 and 120 mJ/cm^2^ fluence prior to the evaluation of their volatiles profile. Overall, aldehyde concentration was not significantly (p > 0.05) affected UV irradiated samples. Primary aldehydes present in untreated samples were 2-methylpropanal, butanal, and acetaldehyde, while in irradiated samples, pentanal, hexanal, and butanal were majorly present. According to Kim and Morr (1996), hexanal can originally be present in the milk while its formation and overall concentration can be increased by the lipid peroxidation caused by the UV light treatment. Similar results were observed in our studies. There was a significant difference (p > 0.05) in overall alcohol concentration in irradiated samples where 1-propanol, n-butanol, and 1-hexanol were major contributors. Among ketones, butane-2,3-dione was majorly present in untreated samples, and its concentration relatively decreased in both irradiated samples (Vazquez-Landaverde et al., 2006). Pereda et al. (2008) reported only one carboxylic acid, i.e., hexanoic acid, in their study because of their less volatility and inefficiency of the system to extract these compounds. In our experiment, we detected seven of them where the major contributor was 2-methyl propanoic acid. Alteration in terpenes was not significant (p > 0.05) in irradiated samples treated with 60 mJ/cm^2^ of UV fluence (Toso et al., 2002). Overall, there was no significant (p > 0.05) difference in overall concentration of aldehydes, ketones, and terpenes of untreated and irradiated samples that received 60 mJ/cm^2^ of UV fluence; however, increase in overall alcohols and esters and decrease in overall carboxylic acids concentration was observed which may be the consequences of light reactions.

**Table 3.**
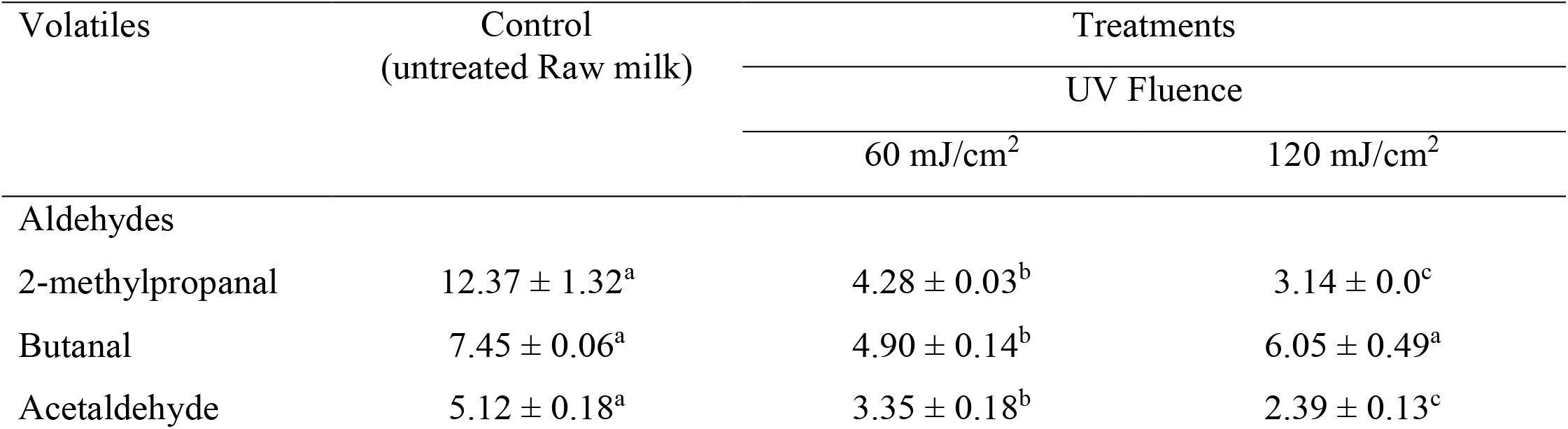
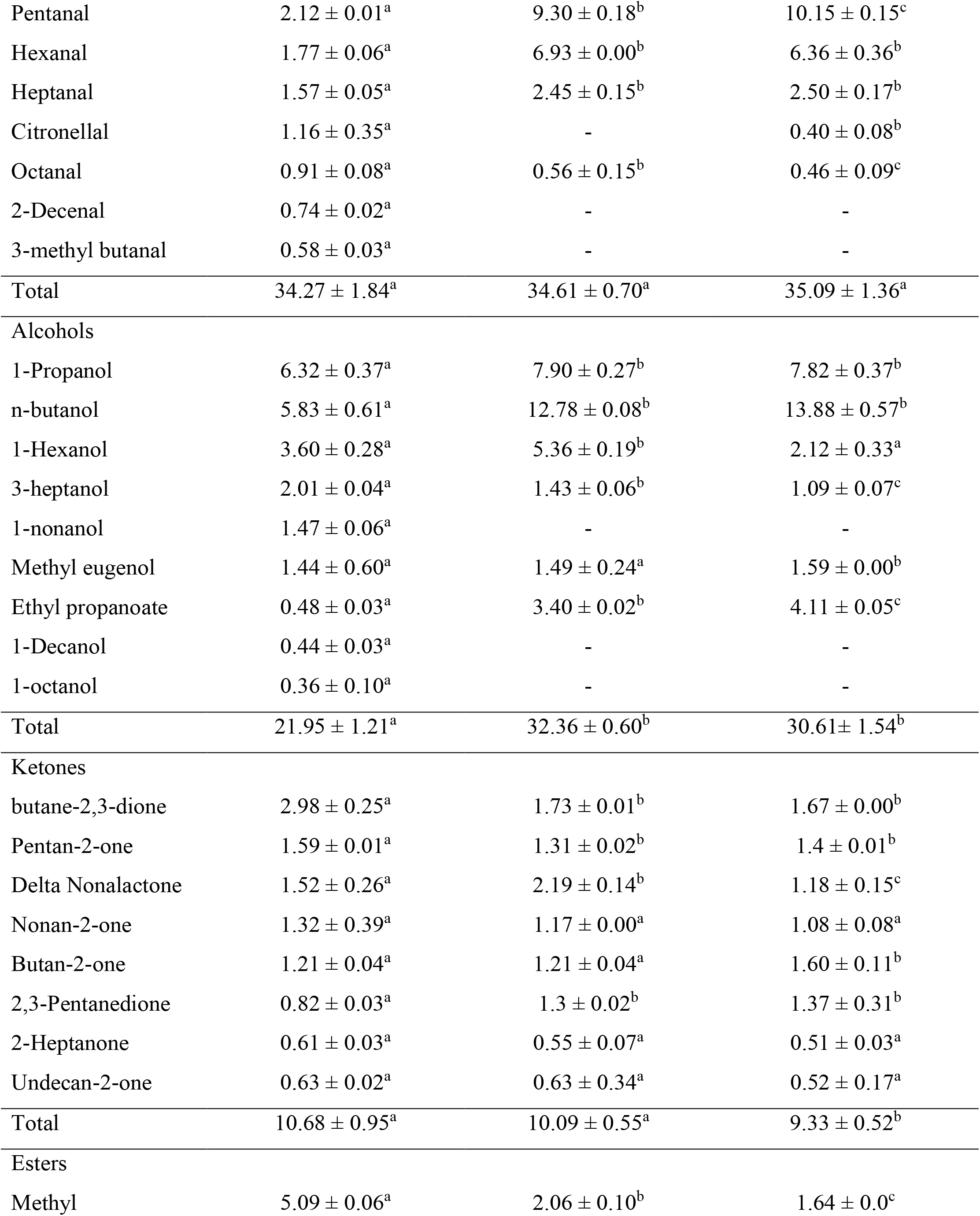
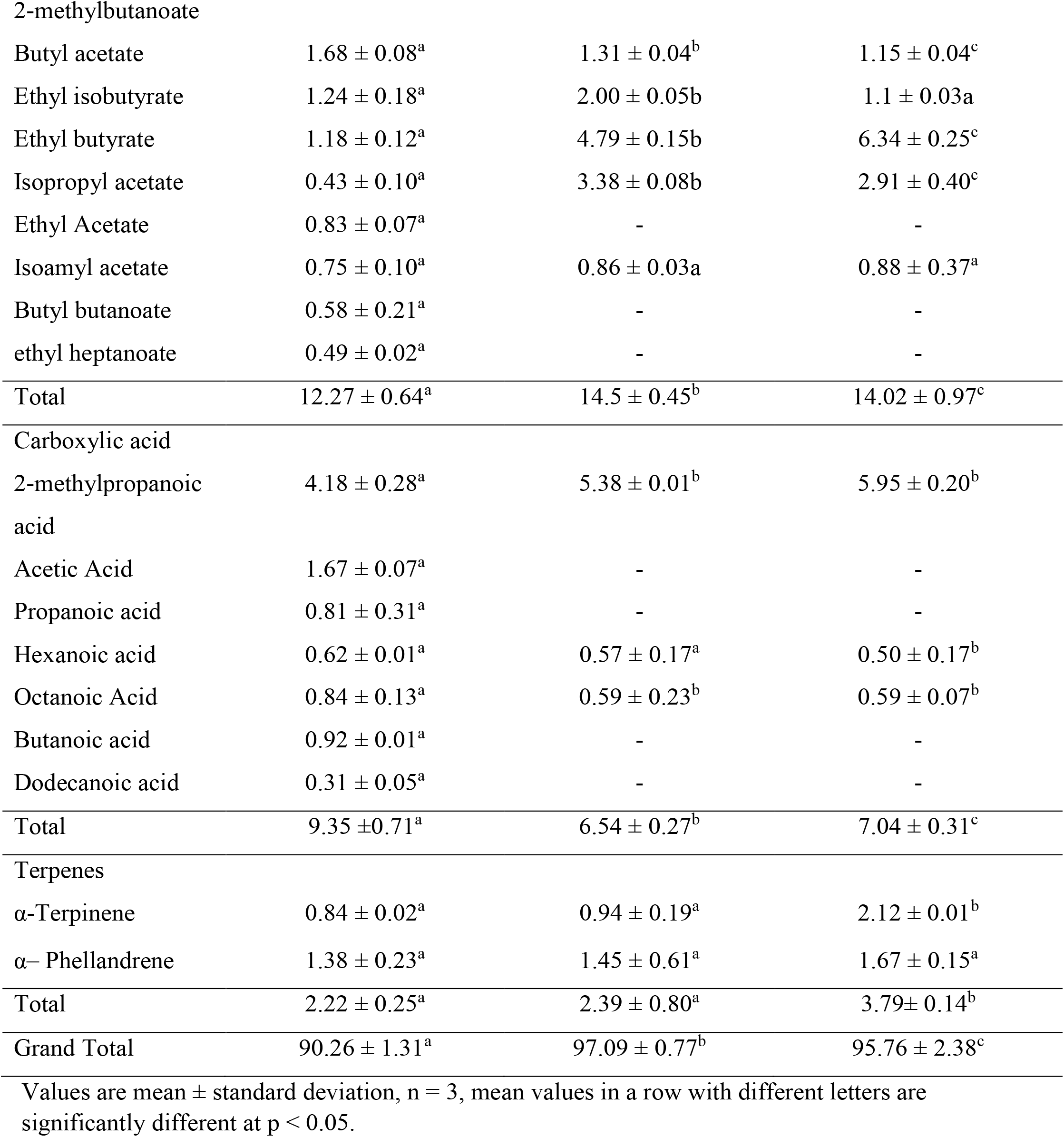
Volatiles profile comparison of the untreated milk samples with irradiated samples treated with UV fluence of 60 and 120 mJ/cm^2^.

### 3.7. Electrical Energy per order (E_EO_) evaluation

The evaluated E_EO_ values for both T1 UV phage and *B. cereus* at 47.55 gph of flow rate are shown in Table 4. T1 UV phage had lower E_EO_ value as compared to *B. cereus* attributed to its lower D_10_ value. Lower E_EO_ value demonstrates the higher efficiency of the system. Since each microorganism has specific UV sensitivity (D_10_ value), E_EO_ value significantly varies with geometric parameters, design of reactor, processing conditions and nature of fluid under testing (Pendyala et al., 2021, Vashisht et al., 2021). E_EO_ value of *B. cereus* was evaluated for each flow rate (Table 4), where it was observed to be decreasing with the increased flow rate attributed to uniform distribution of fluence because of increased turbulence and Dean effect resulted in increased inactivation kinetics. This proves the efficiency of system was more at highest flow rate (47.88 gph) as compared to the lowest one (11.88 gph). At high flow rate two or more reactors are recommended in a series to increase the residence time and hence to deliver the targeted UV fluence. Therefore, a balance needs to be maintained between the chosen flow rate and number of reactors required in a series.

**Table 4.**
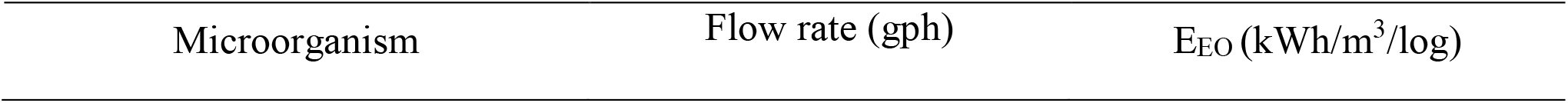
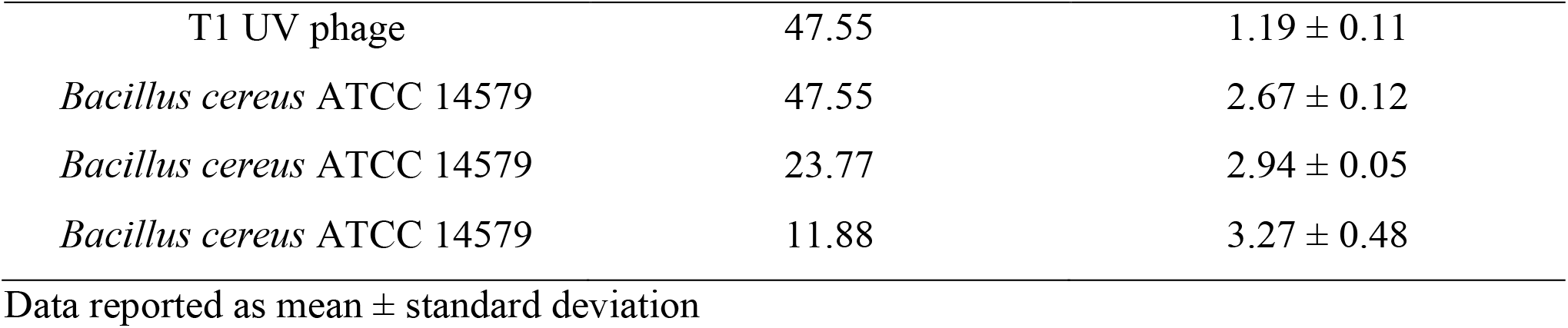
E_EO_ of a Dean flow system to inactivate targeted microorganism under flow rate conditions.

| Microorganism | Flow rate (gph) | $E_{EO}$ (kWh/m<sup>3</sup>/log) |
| --- | --- | --- |
| T1 UV phage | 47.55 | $1.19 \pm 0.11$ |
| *Bacillus cereus* ATCC 14579 | 47.55 | $2.67 \pm 0.12$ |
| *Bacillus cereus* ATCC 14579 | 23.77 | $2.94 \pm 0.05$ |
| *Bacillus cereus* ATCC 14579 | 11.88 | $3.27 \pm 0.48$ |

Data reported as mean ± standard deviation

## 4. Conclusion

A Dean flow UV system was evaluated and experimentally studied using WM as a test fluid due to its high scattering coefficients. Biodosimetry studies were conducted and test results showed that 47.55 gph of flow was the most efficient experimental flow rate for the coiled geometry. Linear inactivation trends were observed for *B. cereus* and T1UV phage demonstrating equivalent fluence during each pass. Lower E_EO_ value also signifies the higher electrical efficiency of the system. Further studies revealed that at 60 mJ/cm^2^ no significant effect on lipid peroxidation and volatiles profile was observed. The reported fluence is enough to inactivate 7.18 log population of *B. cereus* and 12.57 log of T1UV phage. Therefore, it can be concluded that commercial scale UV-C processing of opaque fluids like WM is feasible using an efficient reactor design such as Dean flow system. Further studies on sensory quality of UV exposed WM is warranted in the future.

## Acknowledgments

This project was funded under the Agriculture and Food Research Initiative (Food Safety Challenge Area), USDA, Award numbers; 2018-38821-27732, and 2019-69015-29233. The authors would like to thank Trojan Technologies for providing valuable guidance in this project. **Note:** There are no conflicts to declare.

## CRediT authorship contribution statement

**Pranav Vashisht:** Conceptualization, Methodology, Investigation, Visualization, Writing – original draft. **Brahmaiah Pendyala:** Conceptualization, Methodology, Investigation, Visualization, Writing – original draft, Supervision. **Ankit Patras:** Conceptualization, Methodology, Supervision, Funding acquisition. **Vybhav Vipul Sudhir Gopisetty:** Conceptualization, Methodology. **Ramasamy Ravi:** Methodology, Investigation.

## Notes

### Competing Interest Statement

The authors have declared no competing interest.

